# Structural basis and mechanism of the unfolding-induced activation of an acid response chaperone HdeA

**DOI:** 10.1101/390104

**Authors:** Xing-Chi Yu, Yunfei Hu, Jienv Ding, Hongwei Li, Changwen Jin

## Abstract

The role of protein structural disorder in biological functions is gaining increasing interests in the past decade. The bacterial acid-resistant chaperone HdeA belongs to a group of “conditionally disordered” protein that is activated via an order-to-disorder transition. However, the mechanism for unfolding-induced activation remains unclear due to the lack of experimental information on the unfolded state conformation and the chaperone-client interactions. Here we use advanced solution NMR methods to characterize the activated state conformation of HdeA under acidic condition and identify the client binding sites. The activated HdeA becomes largely disordered and exposes two essential hydrophobic patches of residues for client interactions. The pH-dependent chemical exchange saturation transfer (CEST) result identifies three acid-sensitive regions that act as structural locks during the activation process, revealing a multi-step activation mechanism of HdeA chaperone function at atomic level. The results highlight the role of protein disorder in chaperone function and the self-inhibitory role of ordered structures under non-stress conditions, offering new insights for further understanding the protein structure-function paradigm.

## INTRODUCTION

Intrinsically disordered proteins (IDPs) or proteins containing intrinsically disordered regions (IDRs) constitute about a half of human proteins and are often disease related (1, 2). In prokaryotic proteomes, IDRs are also found to be enriched in proteins involved in pathogenic pathways and essential for invasion of host immune systems (3, 4). Elucidating the functional mechanisms of IDPs or IDRs is critical for understanding the pathogenesis mechanisms of many diseases and also indispensable for a complete elucidation of the protein structure-function paradigm. However, experimental information of disordered proteins is scarce and remains technically challenging to obtain, mainly due to the intrinsic “fuzziness” of the disordered regions. A group of stress-response chaperones in both eubacteria and eukaryotes have been found to adopt well-folded structures under non-stress conditions and become activated via unfolding under stress conditions (5). These include the redox-regulated Hsp33, the temperature-regulated Hsp26, and the acid-activated chaperone HdeA, and are termed “conditionally disordered proteins” (5-10). These proteins represent a unique case of protein disorder, because they undergo an “order-to-disorder” transition during function, which is exactly opposite to the more commonly observed “disorder-to-order” transition either via the “folding-upon-binding” mechanism or by post-translational modifications (11, 12).

The periplasmic chaperone HdeA in enteric bacteria plays a major role during acid stress in protecting a broad range of periplasmic proteins from denaturation-induced aggregation (13-15). The function of HdeA is critical for the survival of pathogenic bacteria when passing through hosts’ stomach, where it interacts with its native clients including the outer membrane proteins (OMPs), as well as chaperones such as SurA and DegP that are essential in the OMP biogenesis pathways (16, 17). HdeA exists as a well-folded homodimer in the inactive state, and its activation requires acid-induced protein unfolding accompanied by dimer dissociation, resulting in the exposure of hydrophobic surfaces for interaction with denatured client proteins (9-10, 13, 18-23). The molecular mechanism of the unfolding-induced HdeA activation is still unclear, and several long-standing questions remain unanswered concerning the following fundamental aspects. First, what is the active conformation of the HdeA chaperone? Second, where is (are) the client binding site(s)? And third, how is HdeA activated and what is the structural basis for such an activation, or more specifically, how does acid induce the exposure of the binding site(s)?

Due to the intrinsic disordered property of activated HdeA, as well as the promiscuous binding between HdeA and client proteins (both are in unfolded or partially unfolded conformations), routine structural studies of the active state conformation and the chaperone-client interactions face severe technical challenges. To address the above issues, we herein employ several state-of-the-art solution NMR techniques that have unique advantages in studying protein disorder and heterogeneous protein-protein interactions at atomic resolution (24-26). The active-state conformation of HdeA has been characterized, which is largely disordered with local residual helical structures. Two hydrophobic segments with relatively extended conformations are identified to play central roles in binding client proteins by using ^19^F NMR method. Further, by using the chemical exchange saturation transfer (CEST) and solvent paramagnetic relaxation enhancement (sPRE) NMR experiments, we observe a pH-regulated dynamic equilibrium between the well-folded inactive conformation and a sparsely-populated partially unfolded conformation. Three pH-sensitive structural regions are identified to function as “structural locks” in regulating the chaperone function activation. Our results provide molecular details of a multi-step activation process of HdeA during acid stress, which demonstrates the central role of disordered regions in the chaperone function and the self-inhibitory role of ordered protein structure.

## RESULTS

### The active state conformation of HdeA

The inactive HdeA is a homodimer under neutral pH conditions, with each monomer consisting four α-helices and α2 locating in the center of the dimeric interface (13, 18, 23). At pH 2, HdeA becomes unfolded and dissociates into monomers, with the estimated *K_d_* value of ~45 μM (13). Under normal sample concentrations for NMR spectroscopy (approximately 100 μM - 1 mM), the activated HdeA is in equilibrium between dimer and monomer conformations, both of which are largely unfolded and show significant signal overlap in the NMR spectra, which greatly hinders spectral analysis (***Supplemental Fig. S1***). In this study, we identified a HdeA-F28W mutant that preserves the chaperone activity (***Fig. 1A-B*** and ***Supplemental Fig. S2A***). This mutant is mainly monomeric at low pH judged by size-exclusion chromatography (***Supplemental Fig. S2B***), and its 2D ^1^H-^15^N HSQC spectrum at pH 1.5 shows a unique set of peaks essentially resembling those corresponding to the monomeric conformation of the activated wild type HdeA (wt-HdeA) (***Supplemental Fig. S3***). Therefore, the HdeA-F28W mutant can represent the monomeric activated state of HdeA, and is used for chemical shift assignments and structural analysis.

**Fig. 1.**
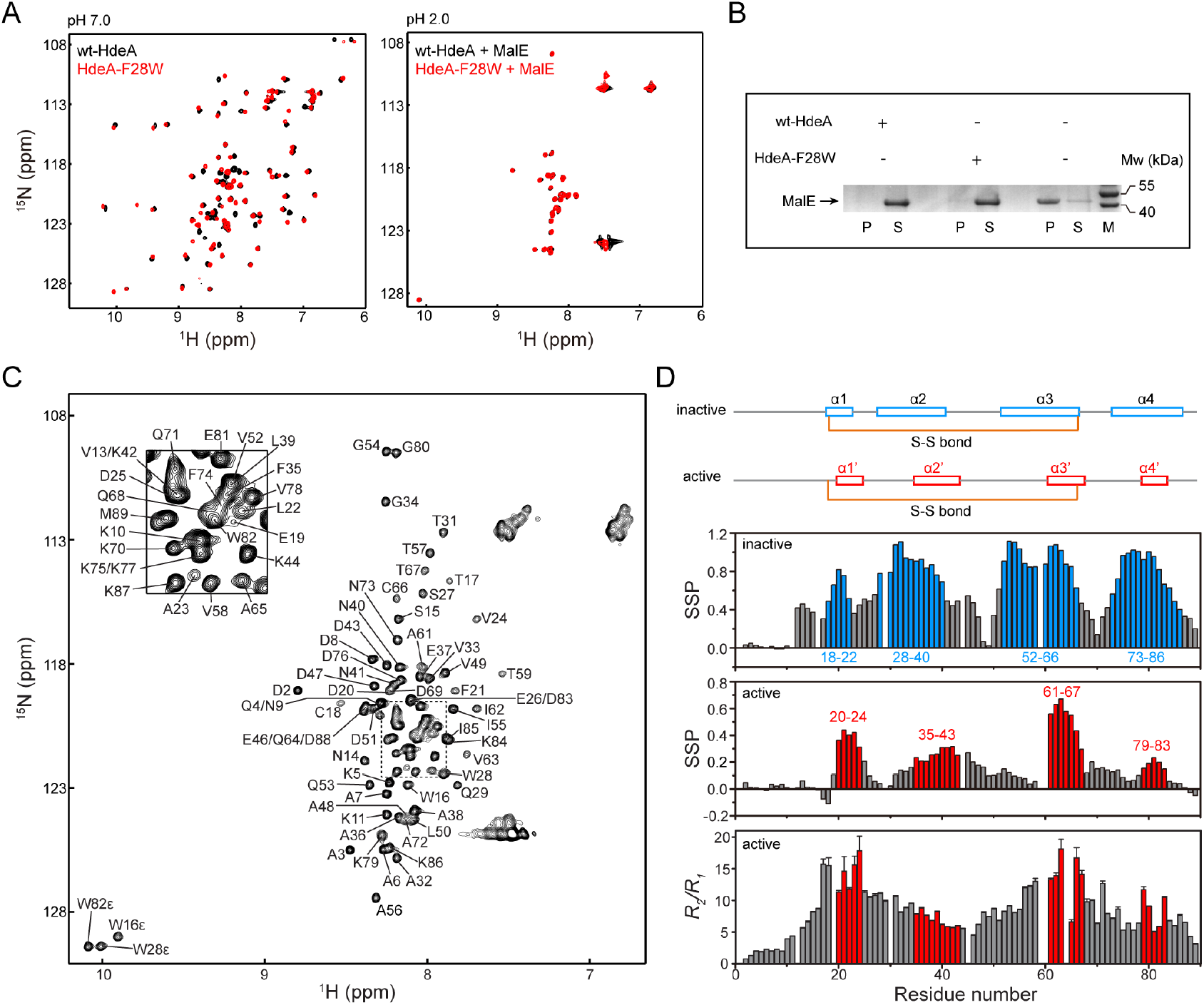
Structural characterization of HdeA in the activated state. (*A*) Overlay of the ^1^H-^15^N HSQC spectra of ^15^N-labeled wt-HdeA (black) and HdeA-F28W mutant (red) at pH 7.0 (left panel), or in the presence of excess unlabeled MalE at pH 2.0 (right panel). (*B*) Chaperone activity of wt-HdeA and HdeA-F28W mutant in protecting the client protein MalE from acid-induced aggregation. The sample of MalE (10 μM) was incubated at pH 1.5 in the presence of equal-molar wt-HdeA (left panel), HdeA-F28W (middle panel) or alone (right panel) for 60 min. P, S and M represent the precipitant, soluble fraction, and protein marker, respectively. (*C*) ^1^H-^15^N HSQC spectrum of HdeA-F28W mutant at pH 1.5 annotated with the backbone resonance assignments. (*D*) Structural analyses of HdeA-F28W at pH 1.5 showing the secondary structure propensity (SSP) scores (middle panel) and the ratios of the backbone ^15^N relaxation transverse and longitudinal rates *R_2_/R_1_* (lower panel), with regions showing propensities of forming secondary structures shown in red. The SSP scores were calculated based on the chemical shifts of ^13^C^α^ and ^13^C^β^ atoms using the program package SSP (27). SSP scores close to 1 indicate high propensity of α-helices. The *R_1_* and *R_2_* relaxation rates were measured on a 700-MHz spectrometer. For comparison, the SSP scores of the inactive HdeA based on the chemical shift assignments of wt-HdeA at pH 3.0 (23) is also shown (upper panel, helical regions shown in blue). The helical elements α1-α4 in the inactive state (blue) and α1’-α4’ in the active state (red) are schematically shown on the top. The intra-molecular disulfide bond is formed between cysteine residues 18 and 66.

By using conventional triple-resonance NMR experiments combined with experiments specifically optimized for disordered proteins, we obtained near complete backbone resonance assignments of HdeA-F28W at pH 1.5 (***Fig. 1C***). Analysis of the secondary structural propensity based on chemical shift information (27) confirms that the protein is largely disordered, whereas four short segments show different tendencies of forming local helical conformations (***Fig. 1D***). These residual helical regions include two segments 20-24 and 61-67, showing close to 40% and 70% helical contents. These two segments are located in the α1-α2 hinge region and the C-terminal half of helix α3 in the inactive structure, and are linked to each other via the strictly conserved disulfide bond between cysteine residues 18 and 66 (***Supplemental Fig. S4***). The other two segments are found in the 35-43 and 79-83 regions, corresponding to the C-terminal half of α2 and the central region of α4, respectively, with the 79-83 regions showing the least helical forming tendency (***Fig. 1D*** and ***Supplemental Fig. S4***). The results are in accordance with the relaxation parameter *R_2_/R_1_* ratios, with higher *R_2_/R_1_* ratios indicating less structural flexibility. The averaged *R_2_/R_1_* value for the segments 20-24 and 61-67 is ~13.6, which is approximately two times of the value of ~7.4 for segments 35-43 and 79-83, and is significantly higher than the N-terminus of the sequence (***Fig. 1D***). This indicates that residual structures are present in the above regions and the local conformation around the disulfide bond maintains the highest rigidity.

### Client-binding sites identified by ^19^F NMR

As demonstrated in our previous study (23), the majority of the backbone amide signals of HdeA in the ^1^H-^15^N HSQC spectra disappear upon binding to client proteins, and the remaining observable resonances correspond to the flexible, charged N- and C-termini (***Fig. 1A***). All signals throughout the 14-72 region are unobservable, making it impossible to identify residues that are most critical for client interactions. To specifically locate the client-binding regions, we used ^19^F NMR spectroscopy to characterize the conformational properties of activated HdeA in both the client-free and -bound states. ^19^F-labeled tryptophan was site-specifically incorporated into 15 different sites of the protein sequence (***Fig. 2A***) and the mutants used for ^19^F-labeling are summarized in ***Table 1***. ^1^H-^15^N HSQC spectra were recorded to confirm that the mutations do not disrupt the overall structure of the inactive state and also do not interfere with client binding (***Supplemental Fig. S5***). Isothermal titration (ITC) experiments (***Supplemental Fig. S6***) and anti-aggregation assays (***Supplemental Fig. S7***) further verified that the mutant retain chaperone activities essentially similar to wild-type HdeA. One-dimensional ^19^F NMR spectra were collected at pH 7.0 and 2.0 for all ^19^F-labeling sites (***Supplemental Fig. S8-S9***), and the results show that the ^19^F chemical shifts are dispersed over a range of ~5 ppm (−47.7 to −52.5 ppm) at pH 7 while clustering around −48.3 ppm at pH 2, which is consistent with acid-induced unfolding (see more details in ***Supplemental Discussion***).

**Fig. 2.**
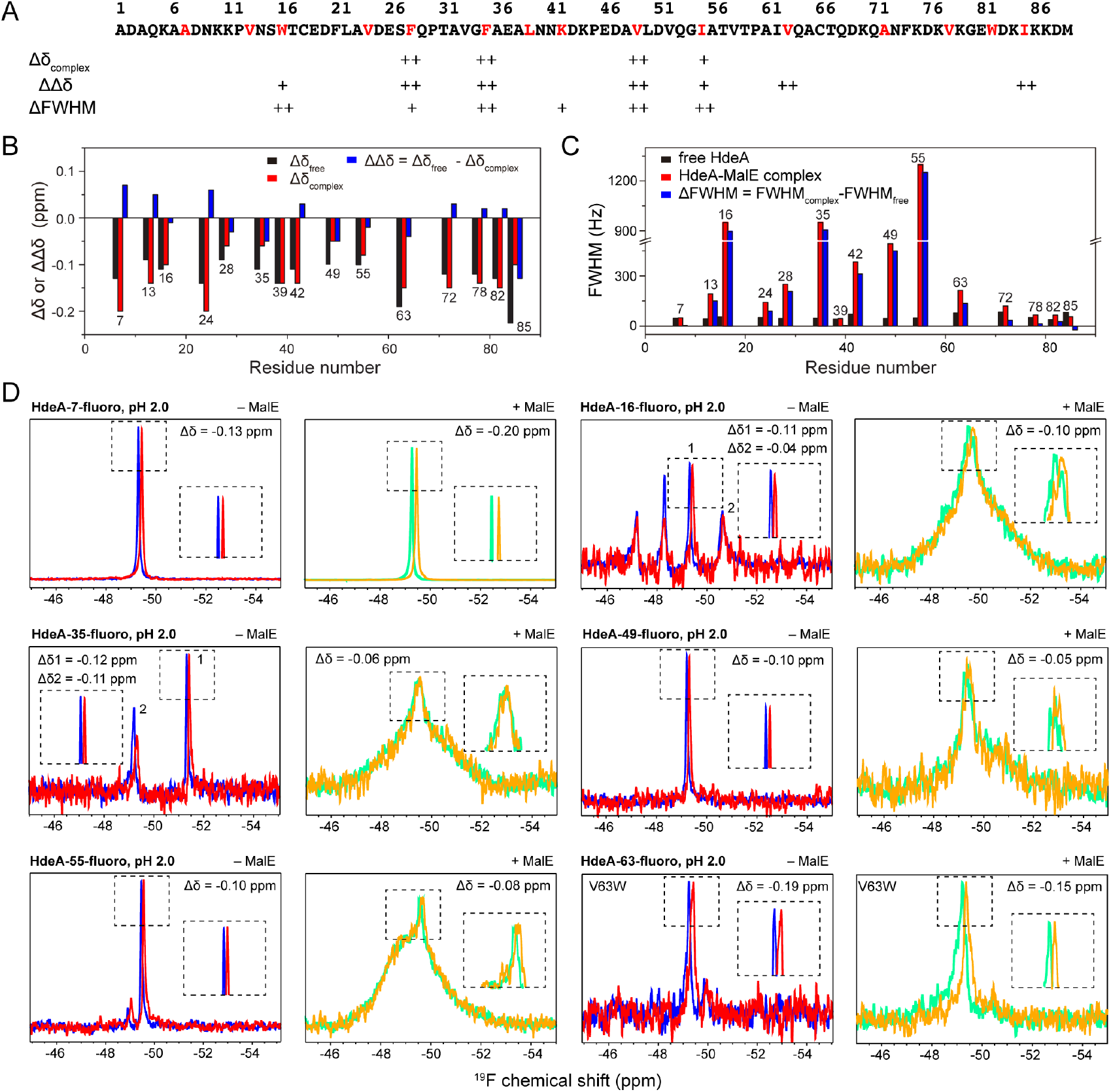
HdeA-client interactions probed by ^19^F NMR. (*A*) The amino acid sequence of wt-HdeA showing the positions for site-specific incorporation of ^19^F probe (colored red). (*B*) The solvent induced isotope shifts Δδ of site-specifically ^19^F-labeled HdeA mutants in the free and complexed state at pH 2.0, and the difference ΔΔδ between the two. The Δδ values are calculated as Δδ = δ(D_2_O) − δ(H_2_O), where δ(D_2_O) is the ^19^F chemical shift measured in 100% D_2_O and δ(H_2_O) is that measured in 90% H_2_O and 10% D_2_O. All Δδ_free_ and Δδ_complex_ values are negative and those closer to zero indicate less solvent exposure in the complexed state and higher possibility of involvement in client binding. The ΔΔδ values are calculated as ΔΔδ = Δδ_free_ − Δδ_complex_, with negative ΔΔδ values indicating less solvent exposure upon client binding. More negative ΔΔδ value indicates larger decrease of solvent exposure upon client binding and thus higher possibility of involvement in client binding. (*C*) The full width at half maximum (FWHM) values of site-specifically ^19^F-labeled HdeA mutants in the free and complexed state at pH 2.0, and the difference ΔFWHM between the two. The ΔFWHM values are calculated as ΔFWHM = FWHM_complex_ − FWHM_free_, with larger ΔFWHM values indicating client binding. The values Δδ_complex_, ΔΔδ and ΔFWHM were independently used as criteria to judge whether the ^19^F-labeled sites are involved in client binding, and the results are indicated below the protein sequence in panel A. ^19^F-labeled sites with Δδ_complex_ values satisfying 0 ppm ≤ Δδ_complex_≤ −0.07 ppm, ΔΔδ values satisfying ΔΔδ ≤ −0.025 ppm, or ΔFWHM values satisfying ΔFWHM ≥ 400 Hz are marked by “++”; those with Δδ_complex_ values satisfying −0.07 ppm < Δδ_complex_ ≤ −0.10 ppm, ΔΔδ values satisfying −0.025 ppm < ΔΔδ < 0 ppm, or ΔFWHM values satisfying 200 Hz ≤ ΔFWHM < 400 Hz are marked by “+”. For the sites 24 and 78, the FWHM values in the free form could not be accurately determined and were estimated by using the averaged value of all other labeling sites (see ***Supplemental Discussions***). (*D*) Representative ^19^F NMR spectra of HdeA mutants with ^19^F probes incorporated at sites 7, 16, 35, 49, 55 and 63 in the free state (left) or in _complex_ with the client protein MalE (right) at pH 2.0. The spectra acquired in a buffer containing 90% H_2_O and 10% D_2_O are shown in blue or green for the free or complexed state, and those acquired in 100% D_2_O are shown in red or yellow for the free or complexed state. Enlarged view of the peaks are shown as in-sets. Sites 16 and 35 show multiple peaks, and the peak corresponding to the unfolded monomeric form (peak 1 as labeled) is used for data analyses (see ***Supplemental Discussions***).

**Table 1.** Summary of HdeA mutants used in the manuscript

| Nomenclature |  | Description |
| --- | --- | --- |
| wt-HdeA |  | wild-type HdeA |
| HdeA-F28W |  | a single mutation with residue Phe28 substituted by tryptophan; mimics the monomeric active state at pH 1.5 |
| HdeA-NΔ9 |  | residues A1-N9 deleted |
| HdeA-F28K |  | a single mutation with residue Phe28 substituted by lysine |
| HdeA-V33K |  | a single mutation with residue Val33 substituted by lysine |
| HdeA-V49K |  | a single mutation with residue Val49 substituted by lysine |
| HdeA-L50K |  | a single mutation with residue Leu50 substituted by lysine |
| Mutants for site-specific <sup>19</sup>F-labeling |  |  |
| <sup>19</sup>F-labeling site | Nomenclature | Mutations |
| 7 | HdeA-7-fluoro | A7W/W16F/W82F |
| 13 | HdeA-13-fluoro | V13W/W16F/W82F |
| 16 | HdeA-16-fluoro | W82F |
| 24 | HdeA-24-fluoro | V24W/W16F/W82F |
| 28 | HdeA-28-fluoro | F28W/W16F/W82F |
| 35 | HdeA-35-fluoro | F35W/W16F/W82F |
| 39 | HdeA-39-fluoro | L39W/W16F/W82F |
| 42 | HdeA-42-fluoro | K42W/W16F/W82F |
| 49 | HdeA-49-fluoro | V49W/W16F/W82F |
| 55 | HdeA-55-fluoro | I55W/W16F/W82F |
| 63 | HdeA-63-fluoro | V63W/W16F/W82F |
| 72 | HdeA-72-fluoro | A72W/W16F/W82F |
| 78 | HdeA-78-fluoro | V78W/W16F/W82F |
| 82 | HdeA-82-fluoro | W16F |
| 85 | HdeA-85-fluoro | I85W/W16F/W82F |

To identify the regions responsible for binding client proteins, we measured the solvent isotope shifts Δδ = δ(D_2_O) − δ(H_2_O) (26) for each labeling site at pH 2.0 in the absence or presence of a native client protein MalE (16) (***Fig. 2B*** and ***Supplemental Fig. S10***). All |Δδ| values are negative and those closer to zero (smaller |Δδ| values) indicate less solvent exposure. Therefore, we expect that the regions of HdeA in direct contact with the client protein to be buried and show the smallest |Δδ| values in the complex sample. In addition, we also expect to see a decrease of solvent exposure (thus negative ΔΔδ values, when ΔΔδ is calculated as Δδ_free_−Δδ_complex_ as shown in ***Fig. 2B***) for residues involved in interaction when comparing the complexed state with the free state. Both criteria unambiguously support the identification of two hydrophobic regions 28-35 and 49-55 as the interaction sites. Residue W16 also appears to contribute to client binding, suggested by the decreased |Δδ| value in the complex state compared to the free state. However, this position exhibits limited solvent exposure in the free state as well, possibly due to the residual secondary structures around the disulfide bond region.

Furthermore, comparison of the signal linewidths in the free and complexed states at pH 2.0 supports the participation of segments 28-35, 49-55 as well as residue 16 in binding client proteins (***Fig. 2C-D***). The significantly increased linewidths of these residues in the presence of the client protein demonstrate direct involvement in binding, whereas the nearly unchanged chemical shifts for all positions reflect the heterogeneous (or promiscuous) nature of the binding. Particularly, the site 55 shows significant line broadening and apparently comprises multiple conformations, indicative of heterogeneous binding (***Fig. 2D***).

Taking all the above factors into consideration, it is clear that the two segments that are rich in hydrophobic residues, namely the 28-35 and 49-55 segments, play central roles in directly binding to unfolded client proteins under acid stress (***Fig. 2A***). This is further supported by the observation that four single mutants HdeA-F28K, HdeA-V33K, HdeA-V49K and HdeA-L50K, each substituting a hydrophobic residue in the two client binding segments by a lysine, show decreased chaperone activity compared to wt-HdeA (***Supplemental Fig. S11*** and *SI Discussion*). In addition, both client binding segments are sandwiched between regions with relatively high propensities of residual helical conformations, but display relatively low propensities themselves (< 20%) of forming helices in the active state (***Fig. 1D***). Notably, the 28-35 segment contributes to the formation of a large portion of the helix α2 in the inactive dimer structure, whereas the V_52_Q_53_G_54_I_55_ tetra-peptide in the 49-55 segment forms the N-terminal tip of helix α3 in the inactive structure. The transition from helical to the more extended random coil conformation may be important for exposure of the hydrophobic residues and thus interaction with client proteins, and the ability of the segments to adopt either helical or random coil conformations may help the chaperone to adapt to a wide variety of substrates.

### pH-regulated order-to-disorder transition

The locations of the 28-35 and 49-55 client-binding sites in the HdeA inactive dimer structure are shown in ***Fig. 3A***. Both segments are solvent-excluded in the dimeric structure and require an order-to-disorder transition to become exposed. In order to elucidate the activation mechanism of HdeA, several molecular dynamics studies have been carried out to explore the conformational properties of unfolded HdeA as well as intermediates along the pH-dependent unfolding pathway (28-32). To provide direct experimental characterizations of such unfolding intermediates, we further used the CEST NMR method that has special advantages in probing sparsely-populated protein conformations (25, 33). Briefly, when the structure of a protein is in exchange between a highly-populated ground state and a sparsely-populated “invisible” (or “excited”) state, saturation on the sparsely-populated state could be transferred to the ground state and result in the signal intensity reduction of the ground state, thus rendering the “invisible” state to become “visible”. The merits of the CEST method include direct measurement of the chemical shifts of the sparsely-populated state, as well as the ability to extract exchange parameters such as the exchange rate *k_ex_* and the relative populations of different conformations.

**Fig. 3.**
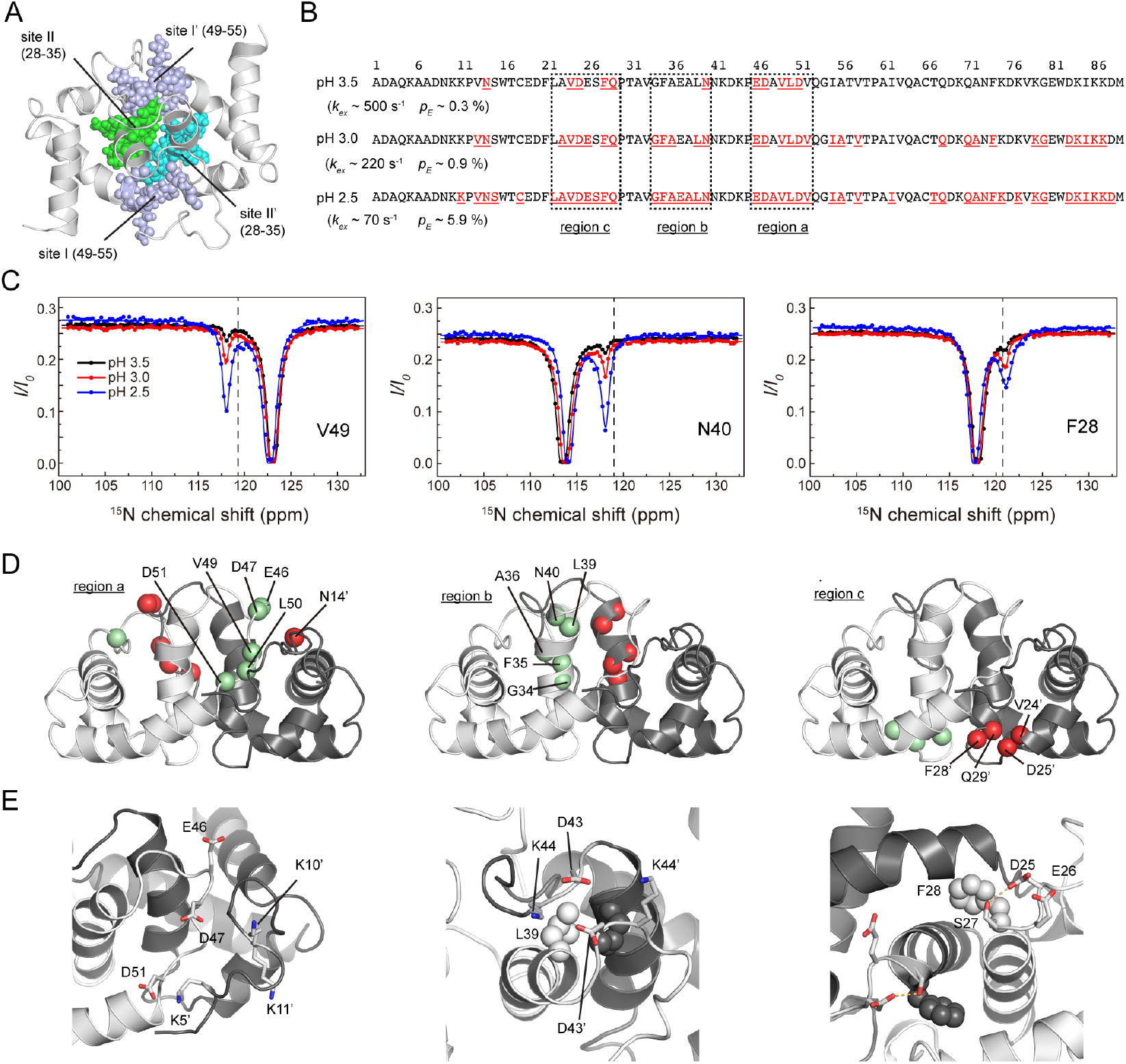
Acid-sensitive structural hot spots identified by CEST NMR. (*A*) Ribbon diagram of the inactive HdeA dimer structure (PDB entry 5WYO) with the client binding sites from both monomers shown as spheres. Binding site I from both monomers are colored in light blue, and the binding site II from the two monomers are colored in green and cyan. (*B*) Residues undergoing conformational exchanges identified by the ^15^N CEST experiments of wt-HdeA at pH 3.5, 3.0 and 2.5 are shown in red in the HdeA protein sequence. Three regions a-c showing sensitive pH responses are depicted by dashed boxes. Exchange parameters *k_ex_* and *p_E_* estimated from global fitting of the CEST data are shown for each pH condition. (*C*) Representative ^15^N CEST profiles of residues in the pH-responsive regions a-c at pH 3.5 (black), 3.0 (red) and 2.5 (blue). The data were obtained using wt-HdeA samples on a 600-MHz spectrometer using a B_1_ field of 14.0 Hz and an irradiation period of duration *T_EX_* = 800 ms. Intensity ratios *I/I_0_* were plotted as a function of position of the weak B_1_ irradiation field, where *I* is the intensity after an irradiation period of duration *T_EX_* and *I_0_* is the intensity in the reference experiment where no B_1_ field was applied. There is a loss of intensity when the weak continuous-wave field is resonant with the major and minor states. The dashed lines indicate the average random coil chemical shift values for the particular amino acid (34). (*D*) Mapping of the pH-responsive residues in regions a-c onto the HdeA dimer structure. Residues from the two monomers are shown in green and red spheres, respectively. Residue N14 spatially close to region a is also shown in the left panel. (*E*) Local conformations of regions a-c in the HdeA dimer structure (PDB entry 5WYO in the left and middle panels and 1DJ8 in the right panel) showing the possible electrostatic or hydrogen bond interactions that contribute to structural stabilization. Charged side chains are shown in sticks, and hydrophobic side chains are shown in spheres. Residues from the second monomer are designated with an apostrophe.

^15^N CEST NMR spectra of the wt-HdeA were systematically recorded at pH values of 4.0, 3.5, 3.0 and 2.5 (***Supplemental Fig. S12***). The experiments detected no excited states for all residues at pH 4.0, whereas sparsely-populated excited conformations were observed for a certain number of residues in the pH range of 3.5 to 2.5, many of which show chemical shifts resembling random coils (34) (***Fig. 3B-C*** and ***Supplemental Fig. S12***). The number of residues harboring the excited state increases as the pH decreases, and a drastic difference was observed between pH conditions of 3.0 and 2.5 (***Fig. 3B***). At pH 2.5 when HdeA is highly activated, over half of the total number of residues show exchanges between folded and unfolded conformations and the residues are spread out in the protein sequence, suggesting that the exchange process corresponds to a global unfolding of the protein structure. This is further supported by the high correlation between the excited state chemical shifts of wt-HdeA at pH 2.5 with the active disordered state chemical shifts obtained from the assignments of HdeA-F28W at pH 1.5 (***Supplemental Fig. S13***). On the other hand, at pH 3.0 and above, order-to-disorder conformational exchanges occur only at local regions and these partially unfolded conformations most probably represent the unfolding intermediates along the pH-induced activation pathway. Global fitting analyses of the CEST data using a two-state exchange model 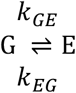 (where G and E stand for the ground and the excited states) further show that the apparent exchange rate constant *k_ex_* (*k_ex_* = *k_GE_* + *k_EG_*) between the folded and unfolded conformations decreases at lower pH, and the relative population *p_E_* of the excited state increases (***Fig. 3B***). The unfolding rate constant *k_GE_* is estimated to be ~1.5, 2.0, 4.1 s^-1^, whereas the folding rate constant *k_EG_* is estimated to be ~500, 218 and 66 s^-1^ at pH 3.5, 3.0 and 2.5, respectively (***Table 2***). The rate constant of unfolding increases of about three-fold when pH decreases from 3.5 to 2.5, whereas the rate constant of refolding decreases of about ten-fold. These results indicate that the energy barrier for the folding-to-unfolding transition decreases along with the decrease of pH, while the energy barrier for the reverse process dramatically increases. This results in a gradual increase of the excited state population, as well as the average lifetime for the excited state, which is ~2.0 ms at pH 3.5, 4.6 ms at pH 3.0 and 15.0 ms at pH 2.5. Moreover, the estimated unfolding rate constant of 4.1 s^-1^ at pH 2.5 is highly consistent with the kinetics parameter (*k* > 3.5 s^-1^) of HdeA unfolding and monomerization reported previously (20), further supporting that the excited states probed by the CEST experiments are on-pathway for HdeA activation.

**Table 2.**
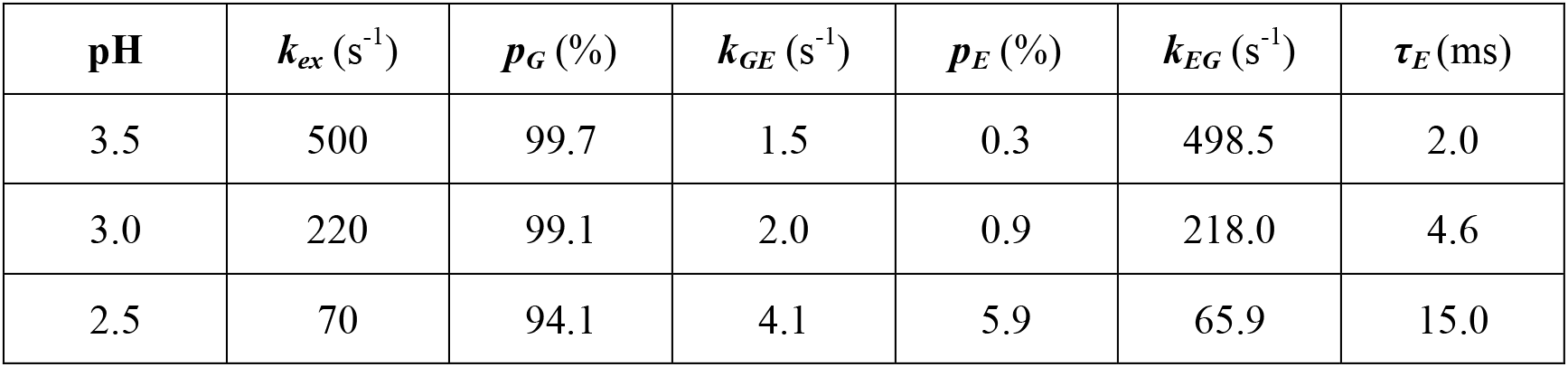
Exchange parameters from ^15^N CEST experiments for wt-HdeA at pH 3.5-2.5

### Conformational transition “hot spots”

By comparing the number and locations of residues showing order-to-disorder exchanges under different pH conditions, we identified three essential acid-sensitive structural “hot spots” in which conformational transitions (dimer dissociation and unfolding) initially occur (***Fig. 3B-E***).

The first region (designated as “region a” hereafter) showing quick acid-response at pH 3.5 is the E_46_D_47_-V_49_L_50_D_51_ segment, which clusters in the C-terminal end of the 50s loop and immediately connects to helix α3 (***Fig. 3C-D***). At pH 3.0, one additional residue V_52_ shows conformational exchanges, whereas at pH 2.5 all residues in the E_46_D_47_A_48_V4_9_L_50_D_51_V_52_ segment undergo exchanges. Notably, the V_49_L_50_D_51_V_52_ tetra-peptide is part of the 49-55 client binding site, and is involved in dimer formation by interacting with the N-terminus of the other HdeA molecule in the homo-dimer via electrostatic interactions (***Fig. 3E***). Protonation of the three acidic residues E_46_, D_47_ and D_51_ could break the electrostatic interactions and lead to local structural loosening. The N14 residue in the N-terminal loop region is spatially close to the E_46_D_47_-V_49_L_50_D_51_ segment in the dimer structure (***Fig. 3D***), and its observed conformational exchanges at pH 3.5 could originate from local collective motions.

The second region (“region b”) corresponds to segment G_34_F_35_A_36_--L_39_N_40_ in the C-terminal end of helix α2 (***Fig. 3C-D***). Compared to region a, this region shows less acid sensitivity since conformational exchanges could be detected for only N40 at pH 3.5. When pH decreases to 3.0, conformational exchanges can be observed for residues G34, F35 and A36 located in the very center of the HdeA dimer. The G_34_F_35_ dipeptide is part of the 28-35 client binding site, and the G34, L39 residues have essential contributions to dimer packing by interacting with G34’, L39’ residues from the other HdeA monomer (here we use an apostrophe to indicate residues from the second HdeA monomer). Notably, residues D43 and K44 in the loop immediately connected to the C-terminal end of α2 show electrostatic interactions with D43’ and K44’ from the other HdeA monomer (***Fig. 3E***), forming an electrostatic “lock” that helps stabilize the dimeric structure. Protonation of D43 could break this “lock” and promote the exposure of the 28-35 client binding site.

The third region (“region c”) corresponds to the V_24_D_25_--F_28_Q_29_ segment at the N-terminal tip of helix α2 (***Fig. 3C-D***). This region shows relatively high acid sensitivity with four residues exhibiting exchanges at pH 3.5, and the number gradually increases with the decrease in pH values. Notably, the F_28_Q_29_ dipeptide is part of the 28-35 client binding site, and the F28 residue contributes to dimer formation by interacting with the P30’ residue. Although the acidic D25 and E26 residues do not show electrostatic contacts with positively charged residues, the carbonyl group of D25 is observed to form a hydrogen bond with the S27 hydroxyl group based on the X-ray crystal structure (***Fig. 3E***). Protonation of the D25 side chain could affect its ability in acting as an electron receptor and destabilizes the local structure.

At pH 3.0 and 2.5, residues in other regions also start to show dynamic exchanges between folded and unfolded states, including the hydrophobic I_55_A_56_-V_58_ segment in the α3 helix which may also participate in client interactions, and the densely-charged C-terminal region harboring the α4 helix (***Fig. 3B***).

### A multi-step activation mechanism of HdeA chaperone function

Based on the above results, a multi-step acid-induced activation mechanism of HdeA chaperone function is derived and schematically summarized in ***Fig. 4***. For clarity, we term the 49-55 segment as client binding site I and the 28-35 segment as client binding site II (***Fig. 3A*** and ***Fig. 4A***).

**Fig. 4.**
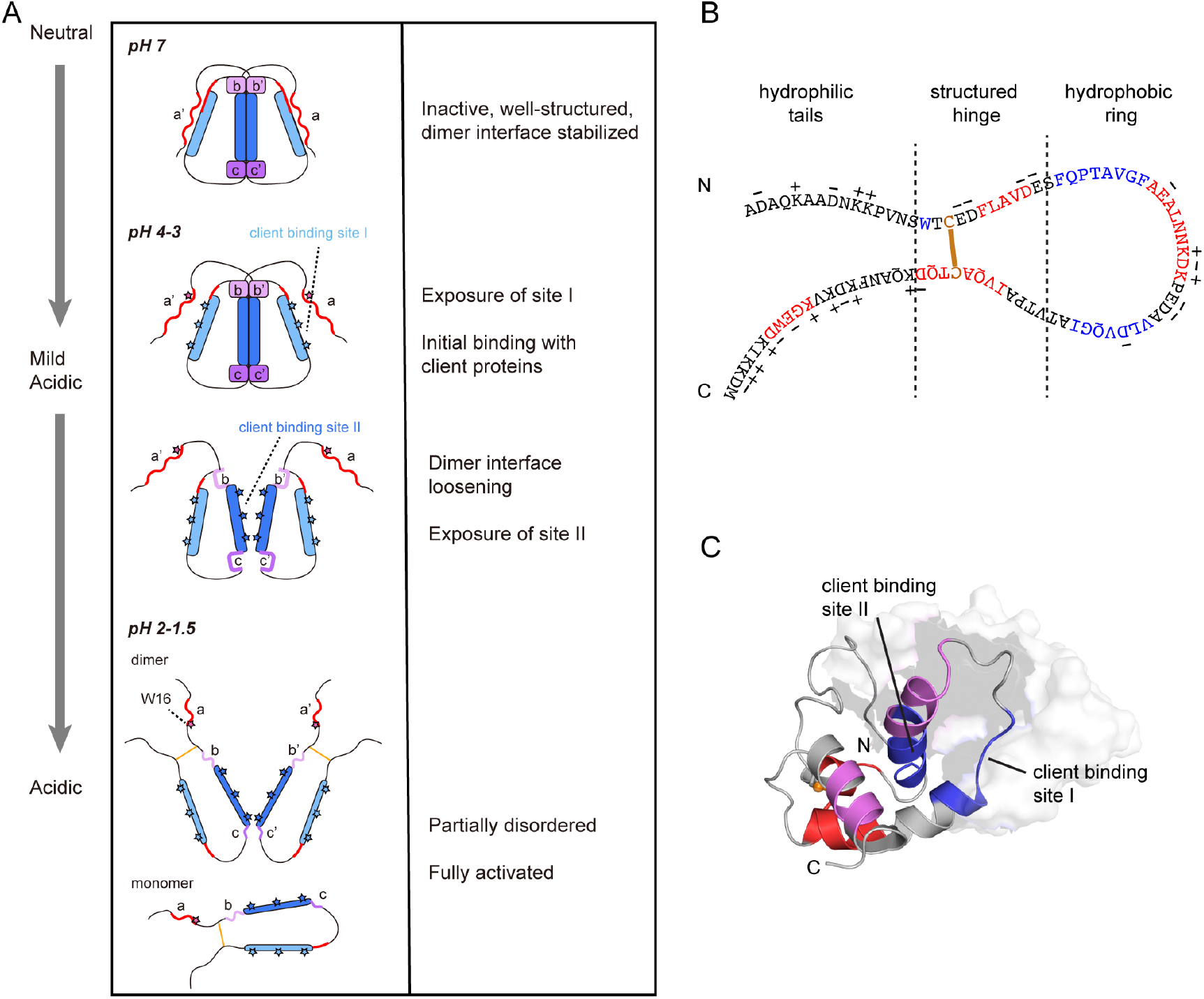
Summary of acid-induced activation mechanism of HdeA. (A) A schematic illustration of the multi-step pH-dependent activation process of HdeA. The two client binding sites I and II are shown in light and deep blue, respectively. As pH decreases, the hydrophobic residues in client binding sites I and II, as well as the W16 residue, gradually become exposed and competent for client binding (illustrated by stars). The pH-sensitive region a and the N-terminal segment that regulate the exposure of site I are colored red, whereas the pH-sensitive regions b and c that regulate the exposure of site II are colored violet and purple, respectively. (B) Illustration The “ring-tail” model of activated HdeA showing the highly charged hydrophilic tails, the relatively structured hinge containing the disulfide bond, and the hydrophobic ring harboring the client binding sites I and II. The client binding sites I and II, as well as residue W16 are colored blue, and segments with relatively high contents of helical conformation are colored red. (C) Mapping of the two client binding sites (colored blue) onto the structure of inactive HdeA dimer (PDB entry 5WYO) showing the proximity of the two binding sites. One monomer is shown in ribbon diagram and the other is shown as the surface representation. The regions showing relatively high helical contents are colored red (highest a-helical forming propensities) and magenta (medium α-helical forming) propensities in the ribbon diagram.

The client binding site I is located at a relative peripheral region in the HdeA dimer structure, and is shielded by the N-terminal segment of the other HdeA molecule. The acidic residues E_46_, D_47_ and D_51_ in the 50s loop have p*Ka* values of 4.07, 4.14 and 3.83 (22) and are in the deprotonated state under neutral and near neutral conditions (pH > 4), ensuring electrostatic interaction with the N-terminal region. As the pH decreases to lower than 4, protonation of these residues results in disruption of the inter-subunit electrostatic contacts and exposure of the client binding site I. The self-inhibitory role of the N-terminal region is supported by the observation that an HdeA-NΔ9 mutant with the N-terminal nine residues deleted show interactions with SurA, another native client of HdeA (13), under elevated pH conditions (***Supplemental Fig. S14A-B***). In addition, the HdeA-NΔ9 mutant also exhibits partial anti-aggregation activity towards the native client protein OppA (13) at pH 4.0, whereas both wt-HdeA and HdeA-F28W are inactive under this pH condition (***Supplemental Fig. S14C*** and ***Supplemental Discussion***). Furthermore, we performed the pH-dependent sPRE experiments, which uses soluble paramagnetic probes to characterize the solvent accessibility of each residue in a protein to provide information concerning protein conformational dynamics. The results show that the inactive HdeA dimer structure is relatively compact at pH 6, with the majority of the residues minimally affected by the paramagnetic probes, whereas residues in the 49-55 segment as well as in the N-terminal region show significantly increased solvent exposure at pH 4 and 3 (***Supplemental* Fig. *S15***). The sPRE results are generally consistent with the NMR hydrogen/deuterium (H/D) exchange data previously reported (22). However, the sPRE data detects a drastic difference of the extent of solvent exposure for the N-terminal residues between pH 6 and 4, which was not observed in the H/D exchange experiments since all resonances from the N-terminal segment already disappeared after buffer exchange even at pH 6. This difference most probably originates from the larger size of the chelated solvent paramagnetic probes compared to water molecules (the probes in the H/D exchange experiments), thus rendering higher sensitivity in observing changes for fast-exchanging amides in the N-terminal segment using the sPRE method.

The client binding site II located in the α2 helix is tightly packed in the structural core of inactive HdeA dimer and could only be exposed via extensive disruption of the dimeric interface. At pH above 4, two acid-sensitive regions (b and c as shown in ***Fig. 3***) present at both ends of the α2 helix stabilizes the dimer interface via inter- and intra-subunit contacts. In addition, previous studies have shown that the protonation of Glu37, an acidic residue showing an exceptionally high pKa value, maximally stabilize the inactive HdeA dimer at pH 5 and protecting residues in the client binding site II from being accessed (10, 22, 28, 35). At pH < 4, acid-induced order-to-disorder conformational exchanges in the b and c acid-sensitive regions loosen the structural “locks” and enables partial dissociation of the α2 helices of the two HdeA molecules. Notably, region b occupies more than half of the total length of the α2 helix, whereas region c locates only at the N-terminal tip of the helix. Therefore, upon acid-induced loosening of these two regions, it appears that the C-terminal part of the α2 helix (the upper part as shown in ***Fig. 3C*** and ***Fig. 4A***) becomes more destabilized. The sPRE data acquired at pH 3-4 show that the solvent accessibility of residues in α2 increases rapidly towards the C-terminal end (***Supplemental Fig. S14***), further suggesting that the C-terminal half is more significantly affected by pH changes and that dimer dissociation is most probably initiated from this end. This scenario is also supported by previously reported H/D exchange data and molecular dynamics simulations (22, 29).

Taken together, the acid-induced activation of HdeA is a multi-step process, which involves the destabilization of three essential “structural locks” that inhibit the chaperone activity under non-stress condition. The initial activation step is the dissociation of the N-terminal loop, which exposes the client binding site I. The subsequent step involves local structural destabilization of helix α2 from both the N- and C-ends, which enables partial exposure of client binding site II. Finally, when pH further decreases, the whole protein structure collapses and becomes fully activated, with both sites exposed for client interactions. In the fully activated state, the N- and C-terminal regions of HdeA form two highly charged, flexible “tails” that help increase the solubility of the HdeA-client complexes, whereas the two essentially hydrophobic client binding sites are held into a “ring” structure by the strictly conserved disulfide bond (***Fig. 4B-C***). Both the spatial proximity of the two client binding sites restricted by the disulfide bond, as well as the higher propensities of adopting random coil conformations for these two segments compared to other regions in the “ring”, could sufficiently increase the effective hydrophobic surface area to facilitate client interactions.

## DISCUSSION

### The self-inhibitory role of the N-terminus

In our previous work, we have hypothesized that the N-terminal charged region plays dual roles in HdeA function, one is to increase the HdeA-client complex solubility, and the other to inhibit the chaperone function under neutral conditions (23). In the current study, we show direct structural evidences that the N-terminus protects one of the two essential client binding sites in HdeA under non-stress conditions, demonstrating its role as a negative regulator of the chaperone activity. The observation that the HdeA-NΔ9 mutant shifts the active pH range to mild acidic conditions strongly supports this hypothesis. Moreover, a previously reported constitutively active HdeA-D20A/D51A double mutant is also consistent with this scenario (10). Based on the solution NMR structure of inactive HdeA dimer (23), we observe that both D51 and D20 play essential roles in stabilizing the N-terminus. D51 forms inter-molecular contact with K5’, whereas D20 forms intra-molecular contact with K11. These two electrostatic interactions, together with the inter-molecular electrostatic contacts between D47 and K10’ described above, act in concert to anchor the N-terminus onto the surface of the client binding site I (***Supplemental Fig. S16***). Disruption of these interactions could result in the release of the N-terminal region and expose site I under non-acidic conditions. However, the HdeA-D20A/D51A mutant shows more severe perturbation on the overall protein folding and its ^15^N-HSQC spectra at pH 7.0 and 5.0 show high content of unfolded conformation (***Supplemental Fig. S17***), which complicates the interpretation of the constitutive chaperone activity at pH > 5 (10). On the other hand, the HdeA-NΔ9 mutant much better preserves the overall folded structure over the pH range of 7.0 to 3.5 compared to HdeA-D20A/D51A (***Supplemental Fig. S17***), while it also exhibits partial chaperone activity at pH 4.0 (***Supplemental Fig. S14***), therefore providing a direct evidence for the self-regulatory role of the N-terminal segment. In the previously study reporting the constitutively active HdeA-D20A/D51A mutant, Foit and co-workers noted that residue D20 is in close proximity to the lysine cluster K10/K11 and interacts with K10 within the same monomer, and interpreted the effect of D20A mutation on HdeA folding as enhancing structural flexibility because of the loss of electrostatic interactions (10). However, due to the lack of electron density for the N-terminus in the crystal structures that were used as the basis of data interpretation, as well as the lack of precise identification of the 49-55 segment as one of the client binding sites, a clear self-inhibitory role of the N-terminal segment could not be established. Moreover, all computational studies reported thus far starts from the crystal structure lacking the first nine residues (10, 28-32, 35), and any local conformational changes involving this segment would have gone unnoticed. Our current results reveal the negative regulatory role of the N-terminal segment and the CEST data can hopefully provide aid in further molecular simulation studies.

Based on our current results, the first step of activation occurs at the client binding site I and involves a simple conformational switch of releasing the N-terminal segment. Thus, without the need for either dimer dissociation or large scale protein unfolding, HdeA can use this patch of hydrophobic residues for initial contact with partially denatured clients under mild acidic conditions (pH 3.5-3). Actually, this can be supported by the appearance of a few N-terminal signals of HdeA (K5, A6 and A7) when it initially interacts with client proteins at pH 3.0-3.8 as reported in our previous study (23), which highlights the release of the N-terminal segment. No new signals from other regions of HdeA were observed upon client interaction, suggesting that the initial contact involves only site I. Apparently, wt-HdeA shows limited chaperone activity in this pH range. However, since the hydrophobic binding area at this binding site may be further extended towards the C-terminal half of helix α3 (e.g. to residue V63) when HdeA becomes globally unfolded at lower pH, the release of the N-terminus and the initial exposure of the 49-55 segment could be important for forming transient complexes at pH > 3 and priming for the formation of more stable complexes upon further unfolding.

More intriguingly, protein sequence alignment of the HdeA and HdeB homologs reveals a nine-residue difference in their N-terminal regions (***Supplemental Fig. S18A***). The HdeB protein is another acid-resistance chaperone but with less well understood functional mechanism. The dimeric structures of HdeA and HdeB are significantly different in terms of dimeric packing, although their monomeric structures are essentially similar (36-39). Recent studies both *in vitro* and *in vivo* suggested that HdeB displays optimal chaperone activity at mild acidic pH values (pH ~4) (17, 38). Up to date, it remains enigmatic as to why HdeB functions at higher pH range when its overall 3D structure remains intact, whereas HdeA is activated only when it is partially unfolded at lower pH values. Sequence comparison between the two homologs reveals that the most conserved residues between these two proteins are clustered in three regions, the 17-24 and 60-74 segments harboring the consensus cysteines, and the 44-58 segment harboring the client binding site I of HdeA (numbered according to HdeA sequence). The sequence conservation in the first two regions may be important for maintaining structural stability, whereas the conservation in the third region implies that HdeB may also utilize the same site for interaction with client proteins. Due to the lack of the N-terminal nine residues in the HdeB sequence, this site is already exposed and competent for client interactions under non-acidic conditions without the need of protein unfolding (***Supplemental Fig. S18B***).

### Unfolding intermediates and chaperone activity

As Foit and co-workers previously suggested, the activation of HdeA chaperone function may not require the protein to be completely unfolded, and that “some of the pH-induced structural changes that accompany HdeA activation are byproducts of acidification rather than an actual requirement for activity” (10). Based on residue hydropathy analysis, the hydrophobic segments in the HdeA primary sequence contain 19-41 and 45-65 and span through helices α1, α2 and α3 (19). Our results show that the client binding sites (28-35 and 49-55) comprise about only half the length of these segments, whereas hydrophobic residues close to the consensus cysteines are not directly responsible for client interactions. The identified binding sites are consistent with previous findings that hydrophobic residues F35, V55 and V58 are essential for client interactions (19, 20). Moreover, the two client binding sites have distinct locations in the HdeA dimer structure and show different pH responses, indicating that HdeA activation follows a pH-dependent step-wise process in which the two sites are activated sequentially and that dimer dissociation (exposure of site II) is more stringently controlled and occurs only at a later stage. Therefore, it is possible that the two sites could act either independently (e.g. at mild acidic conditions when the dimer interface remains intact) or in concert with each other to gain higher chaperone activity (at low pH when HdeA is largely unfolded and the two sites are held in close proximity to provide a larger interaction surface). The sequential activation of HdeA may be important for fine tuning the HdeA-client interactions under different pH conditions.

Studies on HdeA activation mechanism from different groups have convergently pointed to the existence of low-populated partially unfolded intermediates during the acid-induced unfolding (22, 23, 29-31, 35). NMR backbone relaxation measurements have previously suggested that HdeA remains dimeric in the pH range of 6 to 3, however, the hydrogen-deuterium (H/D) exchange data in this pH range indicated that the backbone amides become less protected as the pH decreases (22). In our previous study, we completed the chemical shift assignments of HdeA at pH 3.0, the critical pH value for the order-to-disorder transition, and solved the solution structure which shows a well-folded dimer (23) with no significant changes compared to the crystal structures determined at pH 4 and 3.6 (13, 18). Consistent with this, Salmon and co-workers assigned the ^13^C^α^/^13^C^β^ chemical shifts of HdeA in the pH range of 6.9 to 3.1 and observed modest changes except for acidic residues, indicating that the secondary structures remain relatively unchanged at pH > 3 (35). The apparent discrepancy between preservation of secondary/tertiary structures and the decreased amide protection observed in the H/D exchange experiment (22) led to the proposal of the existence of low-populated partially unfolded intermediates (35), which could also be inferred from the observation that HdeA is able to interact with partially unfolded substrate at pH higher than 3 (23). However, the exact identities of these intermediates and their correlations with chaperone function were poorly understood.

Atomic details of the unfolding intermediates largely rely on computational methods, and a partially unfolded dimeric intermediate (I2) of HdeA was captured in a molecular simulation study showing the characteristics of an almost intact dimeric interface while the helix α4 in one monomer is largely disordered (29). However, the location of the two client binding sites indicates that unfolding and dissociation of helix α4 from the structural core does not play a profound role in exposing the hydrophobic surface responsible for client binding. Inspection of the HdeA dimer structure reveals that the side chains of F74, V78 and W82 from helix α4 form a hydrophobic core within a monomer with F21 from helix α1, V33 from helix α2, and V58, I62 from helix α3, but are distal from either the V49/I55 residues in client binding site I or the bulky aromatic sidechains of F28/F35 in client binding site II (***Supplemental Fig. S19***). Therefore, unfolding of helix α4 within a monomer would not lead to significant exposure of the two hydrophobic binding sites. More importantly, the CEST data indicates that local structure loosening at the three pH-sensitive hot spots starts at a higher pH (pH 3.5) than the unfolding of helix α4 (pH 3.0). Unlike the drastic conformational change associated with helix α4 unfolding, these earlier structural loosening events at the pH-sensitive regions occur in a much more localized manner, but are directly involved in the exposure of client binding sites and thus more correlated with the chaperone activity. Taken together, the unfolding of α4 is more likely to be a byproduct of HdeA structural changes induced by acidification. Nevertheless, unfolding of α4 may contribute to increasing HdeA chaperone activity at low pH by exposing the 58-63 segment in helix α3 as an extension to the client binding site I. Additionally, it is also possible that α4 unfolding could indirectly affect the stability of the dimeric interface and facilitate dimer dissociation.

### Comparison with other chaperone-client interactions

Molecular chaperones are essential for maintaining cellular homeostasis by preventing aggregation of unfolded proteins and assisting protein folding. However, atomic details regarding chaperone-client interactions are poorly understood due to the technical difficulties in obtaining structural data of chaperone-client complexes, which are usually large in molecular sizes and highly dynamic. Recent advances in solution NMR methods greatly enables the structural investigations of these complexes, represented by the atomic-resolution studies of chaperones trigger factor (TF) and SecB in complex with unfolded clients (40, 41). The bacterial periplasmic acid-response chaperone HdeA exhibits some unique characteristics compared to these chaperone systems. A fundamental difference is that while both TF and SecB adopt relatively rigid three-dimensional structures when binding to extended, unfolded clients, HdeA itself is largely disordered and dynamic when carrying out the chaperone function. While SecB forms a disc-like tetramer harboring long hydrophobic grooves that unfolded client wraps around (40), TF binds client proteins in a multivalent mode which is highly dynamic (41). The client binding sites are hydrophobic and spread out on the surface of the two chaperone structures, enabling the maintenance of client proteins in an extended conformation. The two client binding sites in HdeA are hydrophobic, which is a common feature shared among different chaperones. However, the extremely small molecular size of HdeA and the fact that it becomes disordered upon acid-induced activation suggest that the HdeA-client binding mode could be much different from the above two chaperones. The disordered characteristics of the binding sites in HdeA may provide better exposure of the hydrophobic side chains and are favorable for client interactions. The observation that the binding sites show more extended conformations compared to neighboring regions suggests an inverse correlation between the helical-forming propensity and the client-binding activity, and provides a unique demonstration of the link between structure disorder and chaperone function. On the other hand, the client binding sites may adopt transiently formed or sparsely populated helical structures (in particular, residues in site I are estimated to have 10~20% of helical forming propensities), which may fine-tune the local conformation and facilitate promiscuous binding to a broad range of client proteins.

Taken together, our data highlights a complex interplay of both protein structural order and disorder in the regulation of the HdeA chaperone function, in which local structural disorder is responsible for direct binding to client proteins, whereas protein order acts to negatively regulate the chaperone function. To fully understand the functional mechanism of HdeA, atomic-level structural information concerning HdeA-client complexes are essential. The two separated client binding sites and the step-wise activation process suggest the possibility that the mode of HdeA-client interactions could be different under mild acidic and highly acidic conditions, which have potential physiological implications regarding responses to different extent of acid stress, cooperation between HdeA and HdeB homologs, as well as the process of client release (21). However, the disappearance NMR signals in the ^1^H-^15^N HSQC spectra of HdeA upon complex formation most probably reflect the promiscuous and dynamic nature of the binding, and renders atomic-resolution structural determination of HdeA-client complexes through similar approaches as used in the studies of TF and SecB exceptionally difficult. Moreover, much less is known about the conformational states of acid-denatured client proteins and their chaperone-interacting sites, and further investigations focusing on the clients would hopefully shed more light on the HdeA-client interactions.

## Materials and Methods

### Sample preparations

The *E. coli hdeA, malE, surA, oppA* genes and all *hdeA* mutant genes were cloned into pET-28a(+) plasmid (Novagen) and transformed into *E. coli* BL21(DE3)-T1R or BL21 Star(DE3)-T1R strains (Sigma-Aldrich) for protein expression. All protein expression and purification procedures were similar as previously reported (23). The NMR samples were prepared in buffers containing 50 mM sodium phosphate and 45 mM citric acid at different pH conditions.

### Anti-aggregation assay

The chaperone activities of wt-HdeA, the HdeA-F28W mutant and all ^19^F-labeling mutants were tested by the anti-aggregation assay as previously reported (9, 42). MalE was used as the client protein and incubated in a buffer containing 45 mM citric acid, 50 mM sodium phosphate, 150 mM sodium sulfate (pH 1.5) at 25 °C for 60 min with or without the presence of HdeA and its mutants. The addition of 150 mM sodium sulfate was to achieve effective aggregation of MalE at low pH values (43, 44). The MalE concentration was kept at 10 μM, whereas four different concentrations (1, 2.5, 5 and 10 μM) were used for HdeA and its mutants. The presence of MalE in the supernatant or the pellet were analyzed by SDS-PAGE.

For measuring the anti-aggregation activity of the HdeA-NΔ9 mutant at pH 4.0, the client protein OppA was used and the incubation temperature was increased to 35 °C. All other experimental conditions were the same. Control experiments were performed using wt-HdeA and HdeA-F28W at both pH 4.0 and 1.5.

### Chemical shift assignments of the F28W mutant at low pH

For chemical shift assignments of the active monomeric state of HdeA, a sample containing 0.5 mM ^13^C/^15^N-labeled HdeA-F28W mutant was prepared at pH 1.5. NMR spectra were acquired at 25 °C on Bruker Avance 700-MHz spectrometers, equipped with four RF channels and a triple-resonance cryo-probe with pulsed field gradients. Two-dimensional ^15^N-edited heteronuclear single quantum coherence (HSQC) spectroscopy, traditional threedimensional HNCA, HN(CO)CA, HNCACB, HN(CO)CACB, HNCO and HN(CA)CO for protein backbone assignments as well as HNN and HN(C)N experiments especially designed for assigning unfolded proteins were collected (45, 46). All spectra were processed using the software package NMRPipe (47) and analyzed by the program NMRView (48).

### ^19^F labeling

Site-specific incorporation of ^19^F labels into HdeA was achieved by labeling tryptophan residues with 5-fluorotryptophan following published methods (49). Briefly, *E. coli* BL21(DE3) cells harboring plasmids containing the mutant genes were first grown in 1L Luria-Bertani medium at 35 °C. When the OD600 reached 0.8, the cells were collected by centrifugation at 4,000 g and resuspended in 500 mL M9 minimal medium containing with NH_4_Cl and glucose as the nitrogen and carbon sources, 60 mg/L 5-fluoroindole and with 50 mg/L kanamycin. After shaking for 1 hour at 35 °C, protein expression was induced by adding isopropyl-β-D-thiogalactoside (IPTG) to a final concentration of 0.4 mM. Cells were grown for another 8 h and harvested by centrifugation. ^15^NH_4_Cl was used in the M9 media to simultaneously achieve site-specific ^19^F incorporation and uniform ^15^N-labeling.

The wt-HdeA protein sequence contains two tryptophan residues W16 and W82. For ^19^F labeling in these two sites, the HdeA W82F or W16F mutants were used. For ^19^F labeling in all other sites, additional mutations were made based on the HdeA W16F/W82F double mutant (e.g. for ^19^F labeling in position 39, a HdeA W16F/W82F/L39W triple mutant was used). Naming of the site-specific ^19^F-labeled samples follows a “HdeA-xx-fluro” pattern, where “xx” is the number for the amino acid position in the protein primary sequence. A detailed summary of the nomenclature and the corresponding mutations are listed in **Table 1**.

In order to verify that the mutations and ^19^F-labeling do not distort the overall structure of HdeA in the inactive state and do not interfere with client binding, we prepared samples that are site-specifically ^19^F-labeled and simultaneously uniformly ^15^N-labeled, and recorded the ^1^H-^15^N HSQC spectra of the HdeA mutants alone at pH 7.0, 2.0 or in complex with MalE at pH 2.0 (**Fig. S5**). The HdeA-16-fluoro and HdeA-82-fluoro samples using single mutations (W82F and W16F) show the highest spectra similarity to wt-HdeA at pH 7.0, and as expected, other samples use triple mutations and results in larger changes of the chemical shifts in different regions of the protein sequence. Nevertheless, the peak dispersion and homogeneity of the spectra acquired at pH 7.0 indicate that the mutations preserve the overall structure of the inactive HdeA. Furthermore, the HSQC spectra of the HdeA mutants in the presence of the client MalE demonstrate that the mutations do not disrupt HdeA-MalE interactions.

### ^19^F NMR experiments

^19^F NMR experiments were performed at pH 7.0 or 2.0. The preparations for the samples containing ^15^N/^19^F-labeled HdeA mutants alone or in complex with unlabeled substrate MalE at pH 2.0 were similar to previously reported (23). The samples were prepared with 0.5 mM ^15^N/^19^F-labeled HdeA mutants with or without 1.0 mM MalE for the free or complexed states, respectively. Trifluoroacetic acid (TFA) was added to a final concentration of 10 μM as the internal chemical shift reference. The pH conditions of all samples were carefully monitored using methods described previously (23). For all samples, the ^1^H-^15^N HSQC spectra were acquired to ensure that the mutation and labeling did not distort the HdeA conformation or disrupt HdeA-MalE complex formation (**Fig. S5**). One-dimensional ^19^F NMR spectra were acquired on a Bruker 600 MHz spectrometer equipped with a room-temperature BBO probe at 25 °C. A spectral width of 12 kHz and a relaxation delay of 1.5 s were used. A total of 4096 or 100,000 transients were recorded for the free state or HdeA-MalE complex samples, respectively.

For measuring the solvent induced isotope shifts, the samples were initially prepared in a buffer containing 90% H2O and 10% D2O for NMR spectra collection, and subsequently lyophilized and re-dissolved in 100% D_2_O for another round of ^19^F NMR spectra measurement. The solvent induced isotope shifts Δδ values reported in this manuscript are calculated as Δδ = δ(D_2_O) - δ(H_2_O), where δ(D_2_O) is the ^19^F chemical shift measured in 100% D_2_O and δ(H_2_O) is the ^19^F chemical shift measured in 90% H_2_O and 10% D_2_O. The concentration of D_2_O in all samples were carefully controlled. The sample preparations and NMR measurements were repeated to confirm that the ^19^F NMR spectra were well-reproduced. The solvent induced isotope shifts Δδ were also measured at pH 7.0 as a control, where we observed that residues in flexible regions show |Δδ| values in the range of 0.10 to 0.17 ppm, while the |Δδ| value for TFA in 100% D_2_O and in 90% H_2_O/10% D_2_O was 0.14 ppm.

Apart from the solvent isotope shifts, the ^19^F spectra of the HdeA mutants in complex with MalE acquired before and after lyophilization are essentially identical in terms of shape and lined widths, indicating that lyophilization does not change the structure of HdeA in the complex or the mode of interaction between HdeA and the client. This was further verified by the essential similarity between the ^1^H NMR spectra of HdeA-MalE complexes at pH 2.0 before and after lyophilization (the samples were re-dissolved in H2O so that that no hydrogen-deuterium exchange occurs).

### ^19^F NMR data analysis

For analysis of the client binding sites using the solvent induced isotope shifts data, we used two criteria, one is the value Δδ_complex_, which is the solvent induced isotope shift of HdeA in complex with client, the other is the value ΔΔδ, which is the difference between the solvent induced isotope shifts observed in the free and complexes states and is calculated as ΔΔδ = Δδ_free_ Δδ_complex_.

Regarding the value Δδ_complex_, it is expected to be negative and with the maximal absolute value |Δδ_complex_| close to 0.20 ppm (or Δδ_complex_ close to −0.20 ppm) (50). We therefore used the value of 0.20 ppm as an approximation of the largest |Δδ_complex_| expected for a highly solvent exposed site. |Δδ_complex_| values close to 0.20 ppm were observed for sites 7 and 24 in the complexed state, and the average value is ~ 0.12 ppm for all fifteen sites (note that all observed Δδ_complex_ values are negative). During data interpretation, we grouped the 15 sites into three categories based on two thresholds. One is |Δδ_complex_| = 0.10 ppm, which is ~50% of the value for maximal solvent exposure and is also close to the average value of 0.12 ppm. The other is |Δδ_complex_| = 0.07 ppm, which is approximately 30% of the value for maximal solvent exposure. The ^19^F-labeled sites with |Δδ_complex_| value > 0.10 ppm are largely solvent exposed in the HdeA-client complex, those with |Δδ_complex_| value in the range between 0.07 and 0.10 ppm are protected to a certain extent (~30-50% by approximation), and those with |Δδ_complex_| value smaller than 0.07 are considered to be mostly buried in the HdeA-client complex.

Regarding the value ΔΔδ = Δδ_free_-Δδ_complex_, since both Δδ_free_ and Δδ_complex_ are negative, a negative ΔΔδ value means that the site becomes less exposed in the complex state compared to the free state, and therefore indicates possible binding to the client protein. During data interpretation, all residues showing positive or close to zero ΔΔδ values are considered not involved in client binding, and only those showing negative ΔΔδ values are further analyzed. A total of 7 sites (sites 16, 28, 35, 49, 55, 63 and 85) were observed to have negative ΔΔδ, and the most negative data was observed to be −0.05 ppm for sites 28 and 35. We therefore used the value of −0.025 ppm (50% of the most negative value observed) as a cutoff and divided these seven residues into two categories, one satisfying 0 > ΔΔδ > −0.025 ppm and the other with ΔΔδ < −0.025 ppm. The latter group show the largest decrease of solvent exposure upon complex formation and is more likely to be involved in client binding.

For line shape analysis, the ^19^F spectra were processed using the Bruker Topspin 3.5 program with an exponential decaying window function applied to the FIDs during processing. Regarding the value ΔFWHM, which is calculated as AFWHM = FWHM_complex_ − FWHM_free_, the fifteen ^19^F-labeling sites show very large differences, with an average value of ~ 290 Hz and a median of ~ 120 Hz. We grouped the 15 sites into three categories: the first with ΔFWHM < 200 Hz, the second with ΔFWHM in the range of 200-400 Hz, and the third with ΔFWHM larger than 400 Hz. The latter two groups are considered likely to be directly involved in client binding.

### Isothermal calorimetric titration (ITC) experiments

Binding of HdeA or its mutants to the client MalE was measured by ITC using a MicroCal VP-ITC MicroCalorimeter (MicroCal, Northampton, MA) at 25 °C according to the manufacturer’s instructions. All protein samples were dialyzed overnight against a buffer containing 45 mM citric acid and 50 mM sodium phosphate (pH 1.5), and were degassed for 10 min before the titrations. A total of 283-μl concentrated wt-HdeA or its mutants (235 μM) was used as the titrant and added into the MalE solution (1400-ul, 19 μM). The titrations were carried out with a preliminary 2 μl injection (discarded in the data analyses) and followed by 24 injections of 10 μl with an interval of 300 s. The control experiment by titrating HdeA into the buffer was subtracted before data analyses. Note that the buffer used for the ITC measurements does not contain the 150 mM sodium sulfate, which is different from the buffer used in the anti-aggregation assay, so that the MalE protein remains soluble under the experimental condition. All data were analyzed using the program package PEAQ-ITC analysis (Microcal) with different binding models, and the data were best fitted by using the model assuming a single set of unique binding sites with the binding stoichiometry *N* close to 0.5. This reflects a scenario that activated HdeA harbors two binding sites with equal binding affinity, which is consistent with the identification of two binding sites by ^19^F NMR.

### Solvent PRE experiments

Paramagnetic samples were prepared with ^15^N-labeled HdeA (0.6 mM) with paramagnetic probe EDTA-Gd^3+^ (0.5 mM), while the diamagnetic samples were prepared with ^15^N-labeled HdeA alone. An excess of EDTA (1.0 mM) is added in both samples to eliminate the binding of Gd^3+^ ions onto the protein. The ^1^H-^15^N HSQC spectra were collected for both samples at pH conditions of 6.0, 4.0 and 3.0 on a Bruker Avance 800-MHz spectrometer at 25 °C. The spectra of the paramagnetic and diamagnetic samples at each pH condition were carefully compared to ensure that the addition of the EDTA-Gd^3+^ probe did not affect the sample pH and that all signals were well overlaid. The sPRE effects were calculated as the intensity ratio for each residue in the spectra of the paramagnetic and diamagnetic samples. The experimental errors were determined by using duplicated experiments.

### ^15^N CEST measurements and analysis

The ^15^N CEST experiment (33) for wt-HdeA was acquired at 35 °C on a Bruker Avance 600-MHz spectrometer. The sample was prepared in buffers containing 50 mM phosphate, 45 mM citric acid and 10% D_2_O at pH 4.0, 3.5, 3.0 and 2.5 with the protein concentrations of 2.0 mM. A total of 105 2D data sets were acquired with the ^15^N carrier frequencies positioned from 101 ppm to 132.2 ppm at a spacing of 0.3 ppm (18.24 Hz) during the irradiation time of *T_EX_*=800 ms. In all experiments, irradiation field strengths B_1_ of 9.4 ± 0.2 Hz and 14.0 ± 0.2 Hz were used, and a 2.7 kHz field ^1^H decoupling composite pulse sequence (90_x_-240_y_-90_x_) was applied during the *T_EX_* period. Data without using the B_1_ field during the *T_EX_* period was recorded as the reference experiment. B_1_ calibration was carried out following the previously reported methods (51). All the data sets were processed using the NMRPipe program (47), and peak intensities were obtained by NMRView (48). The CEST profiles for the individual residues were generated by calculating the intensity ratios *I/I_0_* versus the varied ^15^N carrier frequencies, where *I_0_* is the intensity measured in the reference spectrum, and *I* is the intensity measured with the application of the B_1_ field. The HSQC spectra of wt-HdeA in the pH range of 4.0-3.0 show a single set of peaks corresponding to the dimeric form as described previously (23), consistent with the estimated micromolar range dimer dissociation constant *K_d_* (13) and the millimolar range protein concentration used to achieve satisfactory signal-to-noise ratio in the CEST NMR experiments. Therefore, the CEST profiles measured under these conditions are recording the conformational exchange processes with the folded dimer as the ground state. Although he HSQC spectra becomes heterogeneous at pH 2.5, only the CEST profiles of resonances corresponding to the dimer species (which could be assigned by tracing the shifts of the signals in the NMR pH titration experiments) were analyzed. Thus, the ^15^N-CEST data at pH 2.5 presented in the current study is also reporting on the conformational exchange processes with the dimer as the ground state.

The CEST data were analyzed using the software package ChemEx (https://github.com/gbouvignies/chemex) or in-house written Matlab scripts from B. Yu and D. Yang (52). For a two-state exchange process 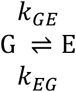, G and E represent the ground state and sparsely populated excited state, respectively. The fractional populations of two states *p_G_* and *p_E_* satisfy the equations *p_G_* = *k_EG_/k_ex_* and *p_E_* = *k_GE_/k_ex_*, with *k_ex_* = *k_GE_* + *k_EG_, p_g_* + *p_E_* = 1, and *p_G_ ≫ p_E_*. The lifetime of the excited state E is given by τ_E_ = 1/*k_EG_*, and the rate constants *k_GE_* and *k_EG_* can be calculated as *k_GE_* = *k_ex_·p_E_*, and *k_EG_* = *k_ex_·p_G_*.

## Acknowledgements

All NMR experiments were performed at the Beijing NMR Center and the NMR facility of National Center for Protein Sciences at Peking University. The authors thank Prof. Zongchao Jia from Queen’s University, Canada for critical reading of the manuscript and helpful suggestions. The authors thank Prof. Zhixin Wang and Prof. Jiawei Wu from Tsinghua University, China for the kind support in the ITC measurements. This research was supported by Grant 2016YFA0501201 from the National Key R&D Program of China to C. J., and Grant 31370718 from the National Natural Science Foundation of China to Y. H.

## REFERENCES

1. Toth-Petroczy A, et al. (2016) Structured States of Disordered Proteins from Genomic Sequences. Cell 167:158–170.

2. Tompa P. (2012) Intrinsically disordered proteins: a 10-year recap. Trends Biochem Sci 7:509–516.

3. Marín M, Uversky VN, Ott T. (2013) Intrinsic disorder in pathogen effectors: protein flexibility as an evolutionary hallmark in a molecular arms race. Plant Cell 25:3153–3157.

4. Gu S, Shevchik VE, Shaw R, Pickersgill RW, Garnett JA. (2017) The role of intrinsic disorder and dynamics in the assembly and function of the type II secretion system. Biochim Biophys Acta 1865:1255–1266.

5. Bardwell JC, Jakob U. (2012) Conditional disorder in chaperone action. Trends Biochem Sci 37:517–525.

6. Reichmann D, et al. (2012) Order out of disorder: working cycle of an intrinsically unfolded chaperone. Cell 148:947–957.

7. Franzmann TM, Menhorn P, Walter S, Buchner J. (2008) Activation of the chaperone Hsp26 is controlled by the rearrangement of its thermosensor domain. Mol Cell 29:207–216.

8. Jaya N, Garcia V, Vierling E. (2009) Substrate binding site flexibility of the small heat shock protein molecular chaperones. Proc Natl Acad Sci USA 106:15604–15609.

9. Hong W, et al. (2005) Periplasmic protein HdeA exhibits chaperone-like activity exclusively within stomach pH range by transforming into disordered conformation. J Biol Chem 280: 27029–27034.

10. Foit L, George JS, Zhang BW, Brooks CL 3^rd^, Bardwell JC. (2013) Chaperone activation by unfolding. Proc Natl Acad Sci USA 110:E1254–1262.

11. Wright PE, Dyson HJ. (2009) Linking folding and binding. Curr Opin Struct Biol 19:31–38.

12. Bah A, Forman-Kay JD. (2016) Modulation of intrinsically disordered protein function by post-translational modifications. J Biol Chem 291:6696–6705.

13. Gajiwala KS, Burley SK. (2000) HDEA, a periplasmic protein that supports acid resistance in pathogenic enteric bacteria. J Mol Biol 295:605–612.

14. Hong W, Wu YE, Fu X, Chang Z. (2012) Chaperone-dependent mechanisms for acid resistance in enteric bacteria. Trends Microbiol 20:328–335.

15. Hingorani KS, Gierasch LM. (2013) How bacteria survive an acid trip. Proc Natl Acad Sci USA 110:5279–5280.

16. Zhang M, et al. (2011) A genetically incorporated crosslinker reveals chaperone cooperation in acid resistance. Nat Chem Biol 7:671–677.

17. Zhang, S, et al. (2016) Comparative proteomics reveal distinct chaperone-client interactions in supporting bacterial acid resistance. Proc Natl Acad Sci USA 113:10872–10877.

18. Yang F, Gustafson KR, Boyd MR, Wlodawer A. (1998) Crystal structure of *Escherichia coli* HdeA. Nat Struct Biol 5:763–764.

19. Wu YE, Hong W, Liu C, Zhang L, Chang Z. (2008) Conserved amphiphilic feature is essential for periplasmic chaperone HdeA to support acid resistance in enteric bacteria. Biochem J 412:389–397.

20. Tapley TL, et al. (2009) Structural plasticity of an acid-activated chaperone allows promiscuous substrate binding. Proc Natl Acad Sci USA 106:5557–5562.

21. Tapley TL, Franzmann TM, Chakraborty S, Jakob U, Bardwell JC. (2010) Protein refolding by pH-triggered chaperone binding and release. Proc Natl Acad Sci USA 107:1071–1076.

22. Garrison MA, Crowhurst KA. (2014) NMR-monitored titration of acid-stress bacterial chaperone HdeA reveals that Asp and Glu charge neutralization produces a loosened dimer structure in preparation for protein unfolding and chaperone activation. Protein Sci 23:167–178.

23. Yu XC, et al. (2017) Characterizations of the interactions between *Escherichia coli* periplasmic chaperone HdeA and its native substrates during acid stress. Biochemistry 56:5748–5757.

24. Burmann BM, Hiller S. (2015) Chaperones and chaperone-substrate complexes: Dynamic playgrounds for NMR spectroscopists. Prog Nucl Magn Reson Spectrosc 86–87:41–64.

25. Sekhar A, Kay LE. (2013) NMR paves the way for atomic level descriptions of sparsely populated, transiently formed biomolecular conformers. Proc Natl Acad Sci USA 110:12867–12874.

26. Kitevski-LeBlanc JL, Prosser RS. (2012) Current applications of ^19^F NMR to studies of protein structure and dynamics. Prog Nucl Magn Reson Spectrosc 62:1–33.

27. Marsh JA, Singh VK, Jia Z, Forman-Kay JD. (2006) Sensitivity of secondary structure propensities to sequence differences between alpha- and gamma-synuclein: Implications for fibrillation. Protein Sci 15:2795–2804.

28. Zhang BW, Brunetti L, Brooks CL 3rd. (2011) Probing pH-dependent dissociation of HdeA dimers. J Am Chem Soc 133:19393–19398.

29. Ahlstrom LS, Dickson A, Brooks CL 3rd. (2013) Binding and folding of the small bacterial chaperone HdeA. J Phys Chem B 117:13219–13225.

30. Ahlstrom LS, Law SM, Dickson A, Brooks CL 3rd. (2015) Multiscale modeling of a conditionally disordered pH-sensing chaperone. J Mol Biol 427:1670–1680.

31. Dickson A, Ahlstrom LS, Brooks CL 3rd. (2016) Coupled folding and binding with 2D window-exchange umbrella sampling. J Comput Chem 37:587–594.

32. Socher E, Sticht H. (2016) Probing the structure of the *Escherichia coli* periplasmic proteins HdeA and YmgD by molecular dynamics simulations. J Phys Chem B 120:11845–11855.

33. Vallurupalli P, Bouvignies G, Kay LE. (2012) Studying “invisible” excited protein states in slow exchange with a major state conformation. J Am Chem Soc 134:8148–8161.

34. Schwarzinger S, Kroon GJ, Foss TR, Wright PE, Dyson HJ. (2000) Random coil chemical shifts in acidic 8 M urea: implementation of random coil shift data in NMRView. J Biomol NMR 18:43–48.

35. Salmon L, et al. (2018) The mechanism of HdeA unfolding and chaperone activation. J Mol Biol 430:33–40.

36. Kern R, Malki A, Abdallah J, Tagourti J, Richarme G. (2007) *Escherichia coli* HdeB is an acid stress chaperone. J Bacteriol 189:603–610.

37. Wang W, et al. (2012) Salt bridges regulate both dimer formation and monomeric flexibility in HdeB and may have a role in periplasmic chaperone function. J Mol Biol 415:538–546.

38. Dahl JU, et al. (2015) HdeB functions as an acid-protective chaperone in bacteria. J Biol Chem 290:65–75.

39. Ding J, Yang C, Niu X, Hu Y, Jin C. (2015) HdeB chaperone activity is coupled to its intrinsic dynamic properties. Sci Rep 5:16856.

40. Saio T, Guan X, Rossi P, Economou A, Kalodimos CG. (2014) Structural basis for protein antiaggregation activity of the trigger factor chaperone. Science 344:1250494.

41. Huang C, Rossi P, Saio T, Kalodimos CG. (2016) Structural basis for the antifolding activity of a molecular chaperone. Nature 537:202–206.

42. Malki, A., Le, H. T., Milles, S., Kern, R., Caldas, T., Abdallah, J., Richarme, G. (2008) Solubilization of protein aggregates by the acid stress chaperones HdeA and HdeB. J Biol Chem 283:13679–13687.

43. Goto, Y., Fink, A. L. (1989) Conformational states of β-lactamase: molten-globule states at acidic and alkaline pH with high salt. Biochemistry 28:945–952.

44. Goto, Y., Takahashi, N., Fink, A. L. (1990) Mechanism of acid-induced folding of proteins. Biochemistry 29:3480–3488.

45. Sattler, M., Schleucher, J., Griesinger, C. (1999) Heteronuclear multidimensional NMR experiments for the structure determination of proteins in solution employing pulsed field gradients. Prog Nucl Magn Reson Spectrosc 34:93–158.

46. Panchal, S. C., Bhavesh, N. S., Hosur, R. V. (2001) Improved 3D triple resonance experiments, HNN and HN(C)N, for HN and ^15^N sequential correlations in (^13^C, ^15^N) labeled proteins: application to unfolded proteins. J Biomol NMR 20:135–147.

47. Delaglio, F., Grzesiek, S., Vuister, G. W., Zhu, G., Pfeifer, J., Bax, A. (1995) NMRPipe: a multidimensional spectral processing system based on UNIX pipes. J Biomol NMR 6:277–293.

48. Johnson, B. A. (2004) Using NMRView to visualize and analyze the NMR spectra of macromolecules. Methods Mol Biol 278:313–352.

49. Crowley, P. B., Kyne, C., Monteith, W. B. (2012) Simple and inexpensive incorporation of ^19^F-tryptophan for protein NMR spectroscopy. Chem Commun (Camb) 48:10681–10683.

50. Kitevski-LeBlanc, J. L., Prosser, R. S. (2012) Current applications of ^19^F NMR to studies of protein structure and dynamics. Prog Nucl Magn Reson Spectrosc 62:1–33.

51. Yuwen, T., Kay, L. E. (2017) Longitudinal relaxation optimized amide ^1^H-CEST experiments for studying slow chemical exchange processes in fully protonated proteins. J Biomol NMR 67:295–307.

52. Yu, B., Yang, D. (2016) Coexistence of multiple minor states of fatty acid binding protein and their functional relevance. Sci Rep 6:34171.

